# Transport and utilization of glycogen breakdown products by *Gardnerella* spp. from the human vaginal microbiome

**DOI:** 10.1101/2022.11.01.514706

**Authors:** Pashupati Bhandari, Janet E. Hill

## Abstract

Multiple *Gardnerella* species frequently co-occur in vaginal microbiomes, and several factors including competition for nutrients such as glycogen could determine their population structure. Although *Gardnerella* spp. can hydrolyze glycogen to produce glucose, maltose, maltotriose and maltotetraose, how these sugars are transported and utilized for growth is unknown. We determined the distribution of genes encoding transporter proteins associated with the uptake of glucose, maltose, and malto-oligosaccharides and maltodextrins among *Gardnerella* species. A total of five different ABC transporters were identified in *Gardnerella* spp. of which MusEFGK_2_I and MalXFGK were conserved across all 15 *Gardnerella* isolates. RafEFGK and TMSP (trehalose, maltose, sucrose and palatinose) operons were specific to *G. vaginalis* while the MalEFG transporter was identified in *G. leopoldii* only. Although no glucose specific sugar-symporters were identified, putative ‘glucose/galactose porters’ and components of a phosphotransferase system were identified. In laboratory experiments, all *Gardnerella* isolates grew more in the presence of glucose, maltose, maltotriose and maltotetraose compared to un-supplemented media. In addition, most isolates (10/15) showed significantly more growth on maltotetraose compared to glucose (Kruskal Wallis, P < 0.05) suggesting their preference for longer chain malto-oligosaccharides. Our findings show that although putative MusEFGK_2_I and MalXFGK transporters are found in all *Gardnerella* spp., some species-specific transporters are also present. Observed distribution of genes encoding transporter systems was consistent with laboratory observations that *Gardnerella* spp. grow better on longer chain malto-oligosaccharides.

**Importance:** Increased abundance of *Gardnerella* spp. is a diagnostic characteristic of bacterial vaginosis, an imbalance in the human vaginal microbiome associated with troubling symptoms and negative reproductive health outcomes, including increased transmission of sexually transmitted infections and preterm birth. Competition for nutrients is likely an important factor in causing dramatic shifts in the vaginal microbial community. *Gardnerella* produces enzymes to digest glycogen, an important nutrient source for vaginal bacteria, but little is known about the mechanisms in *Gardnerella* for uptake of the products of this digestion, or whether *Gardnerella* use some or all of the products. Our results indicate that *Gardnerella* may have evolved to preferentially use a subset of the glycogen breakdown products, which would help them reduce direct competition with some other bacteria in the vagina.

## Introduction

*Gardnerella* spp. are associated with bacterial vaginosis (BV), a dysbiosis characterized by replacement of *Lactobacillus* spp. with a mixture of facultative and anaerobic bacteria from diverse genera such as *Gardnerella, Atopobium, Prevotella, Mobiluncus, Bacteroides*, and others (1). The genus *Gardnerella* is classified into four species; *G. vaginalis, G. swidsinskii, G. piotii* and *G. leopoldii* and nine additional “genome species” have been described (2). *Gardnerella* species differ in their phenotypic characteristics such as *β*-galactosidase production, sialidase activity, and vaginolysin production, which may render some species more pathogenic than others (3, 4).

Multiple *Gardnerella* spp. can co-occur in the vaginal microbiome, however, the relative abundances of the species are different with one usually dominating the mixture. *Gardnerella* spp. in co-culture exhibit scramble competition, which suggests that competition over nutrients is likely an important factor determining the relative abundances of *Gardnerella* spp. in the vaginal microbiome. Glycogen is one important carbon and energy source available for vaginal bacteria. Glycogen accumulates inside the vaginal epithelial cells under the influence of estrogen (5) and is released into the vaginal lumen mainly through the activity of bacterial cytolysins (6). Vaginal glycogen is hydrolyzed into glucose, maltose and malto-oligosaccharides by human and/or bacterial amylases (7, 8) and these products can be utilized by vaginal bacteria including *Gardnerella* spp. to support growth.

Bacteria employ several different transport mechanisms for the uptake of sugars. They can accumulate glucose and other carbohydrates against concentration gradients using ATP (ATP binding cassette (ABC) transporters), ion gradients (major facilitator superfamily (MFS) transporters) or phosphoenolpyruvate (PEP) (PEP-dependent phosphotransferase system, PTS) as energy sources (9). ABC transporters use binding and hydrolysis of ATP to translocate a variety of substrates such as sugars, lipids, drugs, amino acids, etc. across the cell wall (10). They have a characteristic architecture consisting of two transmembrane domains (TMD), two cytoplasmic nucleotide binding domain (NBD) proteins, and membrane anchored substrate binding proteins (SBP) that provide specificity and maintain the direction of transport into the cell. Breakdown products of glycogen are mainly transported via members of the ABC transporter superfamily.

Although all *Gardnerella* spp. can hydrolyze glycogen to produce glucose, maltose, maltotriose and maltotetraose (11), the distribution of transporters for these products, and the extent to which they are used for growth are unknown. Variation in the numbers and types of sugar transporters among species could affect their ability to compete for these sugars. Here, we determined the distribution of genes encoding carbohydrate transporter proteins associated with uptake of glucose, maltose, malto-oligosaccharides and maltodextrins among different species of *Gardnerella*. Furthermore, we measured and compared the growth of *G. vaginalis, G. piotii, G. swidsinskii, G. leopoldii* and *Gardnerella* genome species 3 on glycogen and its breakdown products.

## Methods

### *Gardnerella* isolates

Isolates of *Gardnerella* used in this study were from a previously described culture collection (3) and included a total of 15 *Gardnerella* spp. isolates [three representative isolates each from *G. leopoldii* (GH005, NR017 & VN003), *G. piotii* (GH020, GH007 & VN002), *G. vaginalis* (NR038, NR001 & NR039) and *G. swidsinskii* (NR020, NR021 & NR016), two isolates of *Gardnerella* genome sp. 3 (NR026 & N170) and one from an unknown genome species (NR047, corresponds to the subgroup D based on cpn60 classification system)].

### Whole genome sequencing

Genomic DNA was extracted from all 15 isolates using a modified salting out procedure (12) and quantified using fluorometry (Qubit dsDNA BR assay kit). cpn60 barcode sequencing was performed to confirm the isolate’s identity (13). Briefly, the cpn60 barcode sequence was amplified using primers JH0729 (5’-CGC CAG GGT TTT CCC AGT CAC GAC GAI III GCI GGI GAY GGI ACI ACI AC-3’) and JH0730 (5’-AGC GGA TAA CAA TTT CAC ACA GGA YKI YKI TCI CCR AAI CCI GGI GCY TT-3’) with the following temperature parameters: initial denaturation at 94°C for 5 mins, 40 cycles of (94°C for 30 sec., 50°C for 30 sec. and 72°C for 45 sec.) and final extension at 72°C for 10 minutes. Purified PCR products were sequenced by Sanger sequencing and raw sequence data were analysed to generate the consensus sequence. Finally, each isolate’s identity was confirmed by comparing the consensus sequence to the cpnDB database (14).

Whole genome sequencing libraries were prepared using the SQK-LSK-109 ligation sequencing kit according to the manufacture instructions. Sequencing was performed at Prairie Diagnostic Services (Saskatoon, Canada) on a GridION instrument using a FLO-MIN-106 flow cell. Raw sequences were trimmed to a minimum read length of 1000 base pairs using *filtlong* and trimmed sequences were assembled using *flye* (15). Assembled genomes were annotated using the RAST server (16). The benchmarking universal single-copy orthologues (BUSCO) score (v5.4.3) (17) was used to assess the completeness of the assembled genomes.

### Identification of transporter proteins

Proteome sequences and gene position files for all 15 *Gardnerella* isolates were uploaded to the dbCAN2 webserver (https://bcb.unl.edu/dbCAN2/) (18) and the carbohydrate gene cluster finder (CGC-finder) tool was used to identify carbohydrate gene clusters. CGCs are defined as the genomic regions containing at least one CAZyme gene, one transporter/TC gene and one transcription factor/TF gene (19). CGC-finder performs the carbohydrate transporter annotation of a user sequence by finding its closest hit in the transporter classification database (TCDB), a comprehensive reference database of membrane transport proteins (20). One of the outputs of CGC-finder analysis is a ‘TC prediction output’ file which includes a list of all annotated transporter component proteins along with their TC accession number, an identifier that provides specific information about transporter class, subclass, family, sub-family and predicted substrate(s). Sugar transport proteins in bacteria belong to the carbohydrate uptake transporter-1 family (CUT1) (part of the ABC superfamily), sugar porter family (part of the major facilitator superfamily) and phosphotransferase (PTS) family. All the transporter component proteins assigned to these families were screened to identify transporters associated with uptake of glucose, maltose, malto-oligosaccharides or maltodextrins. The distribution of genes encoding different transporter protein components in all *Gardnerella* isolates was recorded. Gene location information was used to determine the gene organization of the sugar transporters using the SEED viewer (21).

### Phylogenetic analysis of substrate binding proteins

Putative substrate binding protein (MusE and MalX) sequences of MusEFGK_2_I and MalXEFG transporters were aligned using CLUSTALw and neighbour joining consensus trees were built in Geneious Prime version 2022.1.1 (https://www.geneious.com). Trees were visualized using Figtree (V1.4.4).

### Utilization of glycogen and its breakdown products

*Gardnerella* isolates were transferred from -80°C on to Columbia sheep blood agar and incubated at 37°C anaerobically for 48 hours. Isolated colonies were then transferred into 3 ml of NYC III media and incubated anaerobically at 37°C for 48 hours to create a starting inoculum. An aliquot (20 μl) of this starting inoculum was added to 180 μl of fresh modified NYCIII media (mNYC is NYCIII with bovine serum instead of horse serum and without glucose) supplemented with 1% (w/v) of either glucose, maltose, maltotriose, maltotetraose or oyster glycogen in a 96 well plate (in triplicates), and initial OD at 600 nm was measured. The plates were incubated anaerobically at 37°C until the final OD_600_ was taken at 48 hours.

Total growth was determined by subtracting the OD_600_ at 0 h from OD_600_ at 48 h. Planktonic growth was measured by transferring 200 μl of 48 h supernatant from each well into a fresh well and measuring OD_600_. To measure the biofilm formation, a crystal violet staining was performed as described previously (22).

## Results

### Whole genome sequences

Most (12/15) *Gardnerella* genomes were assembled into a single contig while the other three were assembled into two contigs. Coverage estimates ranged from 34× to 569× with an average of 292×. Busco completeness scores were >84% for 13/15 assembled genomes. A full description of read length, coverage and busco score is provided in a Supplementary Table 1. RAST annotation resulted in the identification of 1177-1658 open reading frames per genome (average 1334) (Table 1). These assemblies were used to update the previously published draft assemblies for these isolates (Bioproject number PRJNA394757).

**Table 1:**
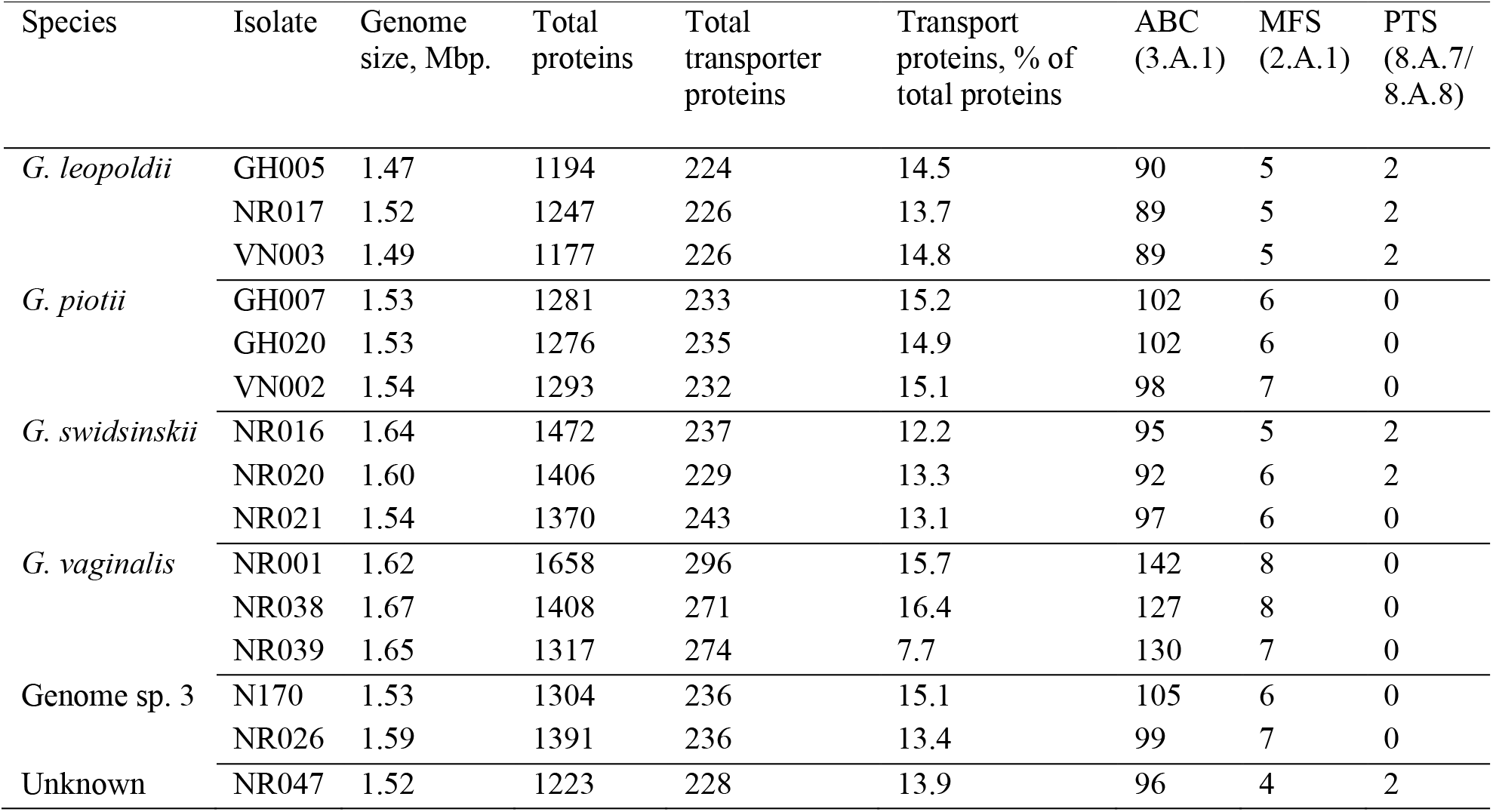
*Gardnerella* isolates used in this study and the distribution of ABC, MFS and PTS transporters

### Distribution of transporter systems in *Gardnerella* isolates

A total of 3,626 transporter proteins were identified among the 20,017 protein sequences from 15 *Gardnerella* isolates uploaded into the dbCAN2 webserver. Of the 3,626 transporter proteins identified, 1,553 (42.8%) belong to the ABC transporter superfamily (TCDB ID 3.A.1), 91 (2.5%) belong to the MFS (2.A.1) and 12 (< 1%) belong to the PTS family (8.A.7/8.A.8). Both the ABC and MFS superfamilies are large and diverse groups of transporter proteins containing many sub-families with different substrate specificities. Within the ABC transporter superfamily, 215/1,553 ABC transporter proteins were assigned to the carbohydrate uptake transporter-1 (CUT1) family (3.A.1.1) with predicted substrate specificity for glucose, maltose, malto-oligosaccharides or maltodextrins. Monosaccharides such as glucose can also be transported via sugar porters that belongs to the sugar porter family (2.A.1.1) within the MFS (2.A.1). Although no proteins belonging to the glucose porter family (2.A.1.1.42) were identified, ‘glucose/galactose porters’ (2.A.1.7.2) (members of the Fucose:H^+^ symporter (FHS) family) were identified in all isolates of *G. vaginalis* and *G. piotii* and two genome sp. 3 isolates but were absent from *G. leopoldii* and *G. swidsinskii*. Proteins corresponding to Enzyme I (8.A.7.1.2) and HPr (8.A.8.1.17) components of the PTS family were identified in all *G. leopoldii*, two isolates of *G. swidsinskii* and one isolate (NR047) from an unknown genome species. In addition, other components of PTS, namely EIIB and EIIC (belong to a PTS-L-ascorbate family (4.A.7.1.4) according to TCDB classification) were identified, however, they were absent in *G. vaginalis* and *G. piotii*. The distribution of different sugar transporter proteins in *Gardnerella* spp. is shown in Table 1.

### Identification of ABC transporters for maltose and malto-oligosaccharides

A total of five different sugar uptake ABC transporter operons were identified in *Gardnerella* spp. MalXFGK (3.A.1.1.27) and MusEFGK_2_I (3.A.1.1.45) operons were conserved across all 15 *Gardnerella* isolates. RafEFGK (3.A.1.1.53) and TMSP (trehalose, maltose, sucrose and palatinose) operons (3.A.1.1.25) were identified only in *G. vaginalis* while the MalEFG transporter (3.A.1.1.44) was identified in *G. leopoldii* only. Out of a total of five sugar specific ABC transporters, four were identified in all *G. vaginalis*, three in *G. leopoldii* and two each in *G. swidsinskii, G. piotii* and others (genome sp. 3) (Figure 1). Genes encoding at least one substrate binding protein and two membrane spanning permease proteins were present in all ABC sugar transporter operons. These operons were unexpectedly lacking genes encoding nucleotide binding domain (NBD) proteins, however, a putative sugar phosphatase (fructose 1,6 biphosphatase (FBP)) was present immediately upstream of the substrate binding protein(s) genes of MusEFGK_2_I and MusE_2_FGK_2_I gene clusters (Figure 1A). Nucleotide binding domain proteins have several characteristic motifs such as Walker A, Walker B, ABC signature, D, H and Q loops (23) but none of these motifs were found in the FBP sequences, suggesting that FBP is not an NDB protein.

**Figure 1:**
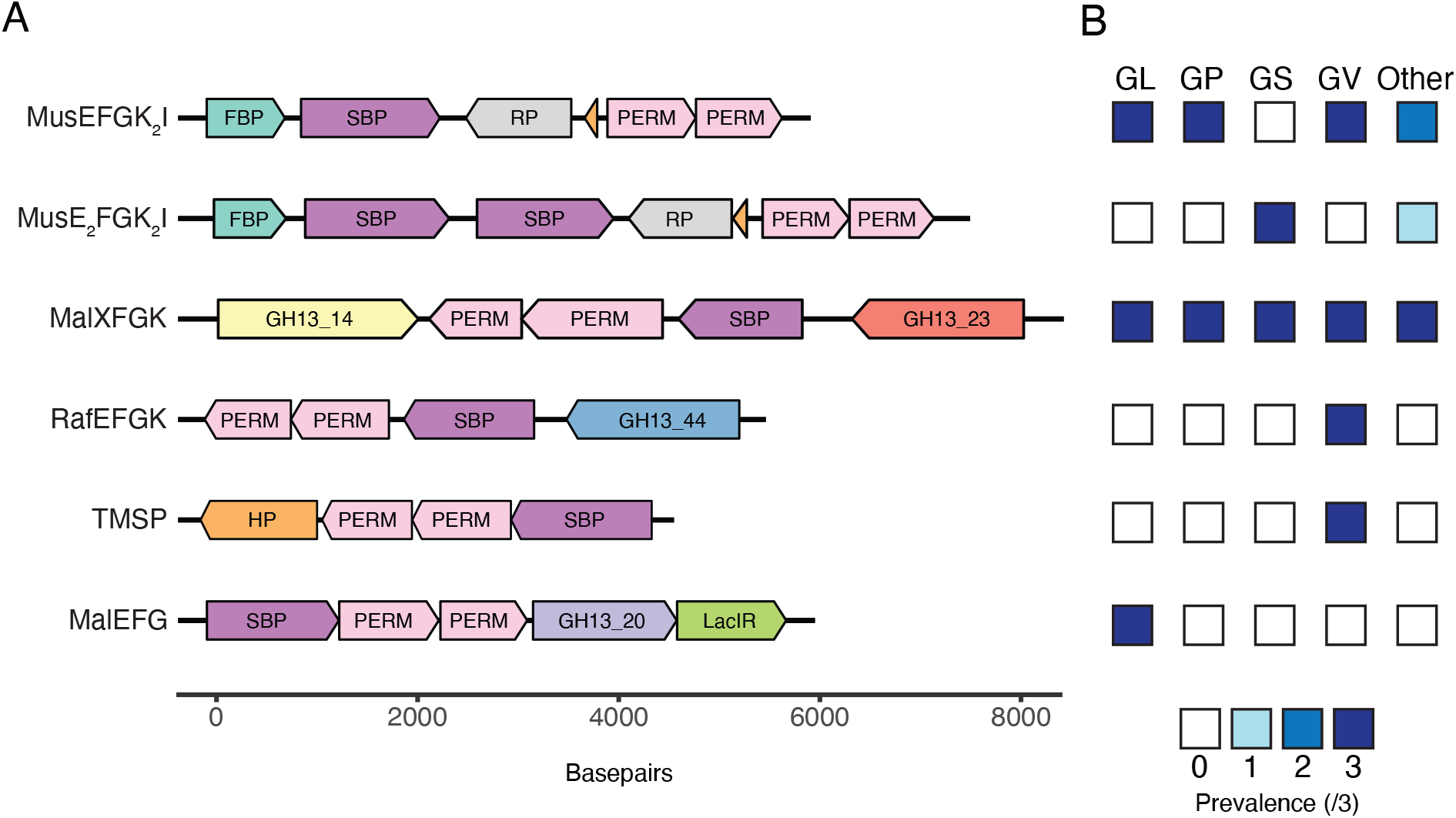
(A) Organization of gene clusters encoding ABC transporters of maltose, malto-oligosaccharides and maltodextrins in *Gardnerella* genomes. Gene clusters identified in representative isolates of *G. vaginalis* NR038 (MusEFGK_2_I, MalXFGK, RafEFGK and TMSP), *G. swidsinskii* NR020 (MusE_2_FGK_2_I) and *G. leopoldii* NR017 (MalEFG) are shown. FBP: Fructose 1,6 biphosphatase, SBP: substrate binding protein, RP: regulatory protein, PERM: permease, GH13_14: pullulanase, GH13_23, GH13_44: *α*-glucosidase, HP: hypothetical protein, GH13_20: *α*-amylase, LacIR: LacI repressor. Hypothetical proteins in the MusEFGK_2_I and MusE_2_FGK_2_I gene clusters are indicated in orange. (B) Prevalence of ABC transporters of maltose, malto-oligosaccharides and maltodextrin in three isolates each of *G. leopoldii* (GL), *G. piotii* (GP), *G. swidsinskii* (GS), *G. vaginalis* (GV) and others (two isolates of genome sp. 3 and one isolate of unknown genome species)).

### Analysis of substrate binding protein sequences from MusEFGK_2_I and MalXFGK

A single gene copy of the gene encoding the MusE SBP was identified in all *G. leopoldii, G. vaginalis, G. piotii* and genome sp. 3 isolates while two copies were identified in all *G. swidsinskii* isolates and one isolate from unknown genome species (NR047) (Figure 1A). MusE SBP sequences annotated by dbCAN2 from 15 *Gardnerella* isolates (n = 18, 2 sequences excluded due to possible frameshifts) were aligned, and the alignment trimmed to a uniform length of 329 amino acids (corresponding to amino acids 157 – 487 of MusE SBP). The phylogenetic tree calculated from this MusE alignment contained two main clusters (Figure 2A). The larger cluster was further divided into two closely related groups of sequences (97-100% identity within each), with some segregation based on species. A second copy of MusE identified in all *G. swidsinskii* and one genome species 3 isolates formed a distinct cluster. Amino acid sequences within this cluster share 77-99% identity. In isolates with two MusE SBP, the paralogs were only 57-61% identical.

**Figure 2:**
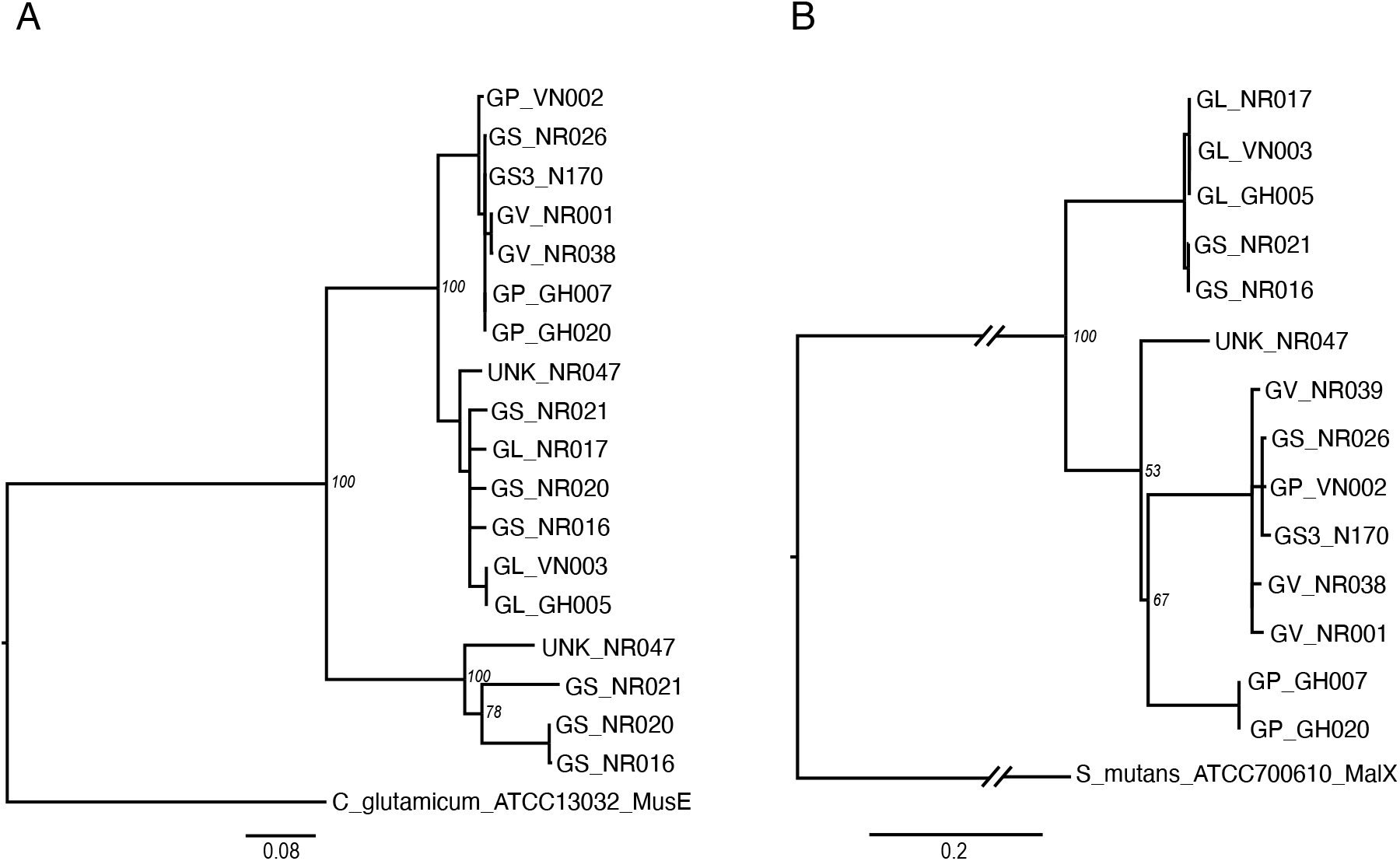
Phylogenetic trees of substrate binding proteins MusE (15 *Gardnerella* isolates) (A) and MalX (14 *Gardnerella* isolates) (B). MusE and MalX trees are rooted with *C. glutamicum* ATCC 13032 and *Streptococcus mutans* ATCC 700610, respectively. Labels indicate species and isolate name (GV, *G. vaginalis*; GP, *G. piotii*; GL, *G. leopoldii*; GS, *G. swidsinskii*; GS3, genome species 3; UNK, unknown species). Trees are the consensus of 100 bootstrap iterations and constructed by Neighbour Joining method using Jukes Canter model. Bootstrap values are shown at the major branchpoints.

A single gene copy of the gene encoding the MalX SBP was identified within the MalXFGK operon of all 15 *Gardnerella* isolates. After removal of one severely truncated sequence, multiple sequence alignments of the remaining 14 sequences trimmed to a uniform length of 268 amino acids (corresponding to amino acids 158 – 425 of MalX SBP) was performed. Three major clusters were formed in the phylogenetic tree of MalX SBP sequences with most sequences segregating according to species (Figure 2B). MalX sequences from all three *G. leopoldii* isolates and 2/3 *G. swidsinskii* isolates formed a single cluster, while sequences from all *G. vaginalis*, both genome sp. 3 and one *G. piotii* clustered separately with good bootstrap support. Sequences within these both clusters were 98-100% identical at the amino acid level. MalX sequences from 2/3 *G. piotii* isolates were clearly distinct from sequences in other clusters with 100% bootstrap support while the MalX SBP of NR047 (unknown genome species) did not cluster with any other MalX sequences.

### Utilization of glycogen and its breakdown products

All *Gardnerella* isolates attained higher OD_600_ values in mNYC supplemented with glucose, maltose, maltotriose, maltotetraose or glycogen compared to the basal media demonstrating their ability to utilize glycogen or its breakdown products for growth (Figure 3). In addition, more growth was observed on maltotriose and maltotetraose compared to glucose or maltose in all isolates suggesting their preference for the longer chain malto-oligosaccharides. Most isolates showed significantly higher growth in the presence of maltotriose (12/15), maltotetraose (15/15) and glycogen (12/15) compared to mNYC and 10/15 isolates showed significantly higher growth in the presence of maltotetraose compared to glucose (Kruskal Wallis test, *P <* 0.05) (Figure 3). To determine the contribution of biofilm to 48 h absorbance, planktonic growth, total growth and biofilm formation were measured for three selected *Gardnerella* isolates. Negligible amounts of biofilm were detected by crystal violet staining, demonstrating that the isolates grew primarily in planktonic mode (Supplementary Figure 1). This was not surprising as in our experience, *Gardnerella* spp. do not produce biofilm in serum containing media and the modified NYC media used contains fetal bovine serum.

**Figure 3:**
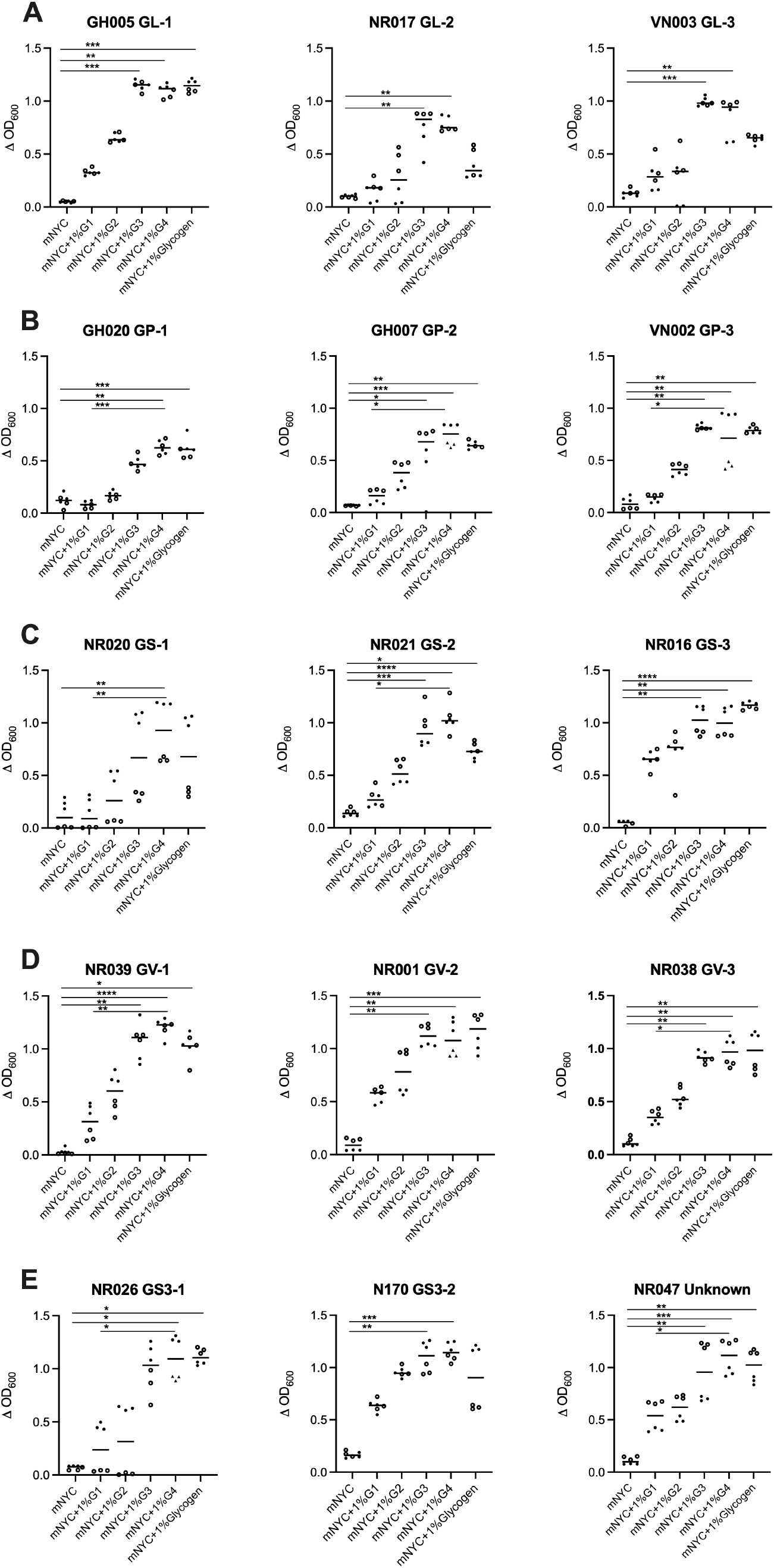
Growth of 15 *Gardnerella* isolates in mNYC III media, mNYC III supplemented with 1% glucose, 1% maltose, 1% maltotriose, 1% maltotetraose or 1% glycogen. Panel A: *G. leopoldii*, B: *G. piotii*, C: *G. vaginalis*, D: *G. swidsinskii* and E: Two isolates from *Gardnerella* genome sp. 3 and one unknown genome species. Data from two independent experiments (indicated by different shapes) each with three technical replicates are shown. *P* values (Kruskal Wallis test) of <0.05 (*), <0.01 (**) and <0.001 (***) are indicated. Horizontal lines indicate median.

## Discussion

Glycogen is one important carbon and energy source for vaginal microbiota. In the vagina, glycogen released from epithelial cells is hydrolyzed by human and/or bacterial amylases into mainly glucose, maltose, maltotriose and maltotetraose (7, 8), providing nutrients for the microbiota. Bacteria are equipped with several transporters with different and sometimes overlapping substrate specificities to transport mono-, di- and trisaccharides (9) and the transport capabilities of these systems can greatly influence the ability of bacteria to compete for available sugars. Although *Gardnerella* spp. can release glucose, maltose, maltotriose and maltotetraose from glycogen (11) the ability of each species to transport and utilize these sugars has not previously been investigated. Competition for glycogen breakdown products could play an important role in determining the relative abundances of *Gardnerella* spp. in the vaginal microbiome.

Out of five different ABC sugar transporters identified, only MusEFGK_2_I and MalXFGK were conserved across all *Gardnerella* isolates included in this study. The maltose uptake system (MUS) belongs to the ABC transporter class (3.A.1.1.45) and has been well characterized in *Corynebacterium glutamicum*. This transporter is encoded by the MusEFGK_2_I operon and transports mainly maltose and maltotriose (25). In all *Gardnerella* isolates, the MusEFGK_2_I gene operons were found to encode at least one SBP and two permeases except *G. swidsinskii* and one isolate from an unknown genome species where two gene copies were identified. Multiple copies of SBP are common in ABC transporters, which can enhance transporter capacity and broaden substrate specificity (26). In our current study, where two MusE SBP genes were identified, the encoded protein sequences were divergent (57-61% identical) and clustered separately in the phylogenetic analysis (Figure 2A), which likely indicates different substrate specificity.

The MalXFGK transporter (3.A.1.1.27) has been characterized in *Streptococcus mutans* and is involved in the transport of maltose, maltotriose, malto-oligosaccharides (DP≤7) and maltodextrins (27). In *Gardnerella*, the MalXFGK operon also encodes an *α*-glucosidase and a pullulanase (Figure 1A). This intracellular *α*-glucosidase has been previously shown to hydrolyze *α*-1,4 glycosidic bonds of malto-oligosaccharides to release glucose (28) while pullulanase is a debranching enzyme that breaks down *α*-1,6 glycosidic bonds (29). Thus the MalXFGK operon encodes all of the components needed to import malto-oligosaccharides and/or maltodextrins for debranching and hydrolysis by pullulanase and *α*-glucosidase to release glucose. Interestingly, the MalX SBP sequences we identified (Figure 2B) were more diverse than MalE sequences from the same isolates, raising the possibility that substrate specificity of the MalXFGK transporters may differ among *Gardnerella* spp.

In contrast to the conserved MusEFGK_2_I and MalXFGK operons, RafEFGK (3.A.1.1.27) and TMSP (3.A.1.1.25) were present only in *G. vaginalis* while MalEFG (3.A.1.1.44) was identified only in *G. leopoldii* (Figure 1B). RafEFGK is involved in the transport of *α*-1,6 linked glucosides and galactosides such as raffinose, panose, stychose, melibiose, and isomaltose. Similarly, TMSP is responsible for the uptake of disaccharides mainly trehalose, maltose, sucrose and palatinose while MalEFG is associated with uptake of maltose and maltodextrins (20). Overall, *G. vaginalis* has the highest number and diversity (4 of the 5 ABC transporters identified) of sugar specific ABC transporters, which is consistent with a previous finding that *G. vaginalis* encodes significantly more proteins associated with carbohydrate transport and metabolism compared to others (30).

ATPase components of the ABC transporters are required to energize the substrate transport across the membranes, however they do not confer the specificity (31). All 5 sugar ABC transporter operons identified in *Gardnerella* spp. lacked an ATPase protein gene. The fulfillment of this role by a protein encoded elsewhere in the genome is certainly a possibility. An ATPase encoded by a distant locus serves as an NBD and energizes the transporters involved in uptake of maltodextrin in *B. subtilis* (32) and *S. pneumoniae* (33). In addition, the ATPase domain of one ABC transporter can interact with an alternative ABC transporter complex to facilitate the sugar transport (27). The identification of NBD proteins that fuels ABC transport in *Gardnerella* spp. requires further experimental investigation.

Transport of monosaccharides such as glucose against concentration gradients is facilitated by porters such as symporters, uniporters or antiporters of the MFS type (9). MFS transporters exhibit specificity for sugars, drugs, amino acids, peptides, polyols, nucleosides, organic and inorganic ions and many other solutes (20). Although members of the glucose porter family (2.A.1.1.42) were absent in all *Gardnerella* isolates we examined in the current study, a putative ‘glucose/galactose’ porter was identified in all isolates of *G. piotii, G. vaginalis* and two isolates of genome sp. 3. This putative ‘glucose/galactose’ porter has been reported in *Brucella abortus* (34) (a sequence with 33% amino acid identity to the *Gardnerella* glucose/galactose porter), however, its actual function has yet to be demonstrated. The uptake of glucose and other monosaccharides can also be catalyzed by PEP-dependent PTS. PEP serves as an energy source and phosphoryl donor which is transferred via different cytoplasmic proteins (EI, HPr and EIIA) to the transported sugar bound to membrane components (EIIB and EIIC) of the PTS (35). Putative EI (8.A.7.1.2), HPr (8.A.8.1.17) and EIIBC (4.A.7.1.4) components were identified in all isolates of *G. leopoldii* and 2/3 isolates of *G. swidsinskii*, however they were absent in *G. piotii* and *G. vaginalis*. It is interesting to note that the putative ‘glucose/galactose’ porter and PEP-PTS are mutually exclusive in the study isolates (Figure 1B). Our observations that *G. piotii, G. vaginalis* and genome sp. 3 all grew more on glucose supplemented media compared to un-supplemented media suggests that the ‘glucose/galactose’ porter provides glucose transport in these species, while in other *Gardnerella* species it occurs via the PEP-PTS transporter.

The observed distribution of transporter systems with predicted specificity for glycogen breakdown products among *Gardnerella* spp. is consistent with their ability to use all these substrates for growth (Figure 3). The fact that all *Gardnerella* isolates exhibited more growth in mNYC supplemented with maltotriose and maltotetraose compared to glucose and maltose supplemented media further suggests that the efficiency and capacity of transporter systems for glucose and maltose may be limited compared to those dedicated to longer chain oligosaccharides. In contrast, vaginal lactobacilli are reported to grow better in glucose and maltose compared to maltodextrins (7, 36). Together these findings suggest that *Gardnerella* and *Lactobacillus* spp. may occupy different nutritional niches in the vaginal microbiome. Further experiments are required to confirm the sugar preference of *Gardnerella* and *Lactobacillus* spp. and to determine the extent to which they compete for these substrates.

Our findings show that putative MusEFGK_2_I and MalXFGK transporters are conserved across all *Gardnerella* isolates and all *Gardnerella* spp. grow better in presence of malto-oligosaccharides compared to glucose and maltose. Although RafEFGK and TMSP transporters were specific to *G. vaginalis* and *G. leopoldii*, respectively, whether they confer competitive advantage for these species in vaginal microbiome by selective sugar uptake remains to be determined. Taken together, our results show that utilization of common energy sources is an important consideration in understanding the forces at work in initiating and maintaining dysbiotic states in the vaginal microbiome.

## Acknowledgements

This research was supported by a Natural Sciences and Engineering Research Council of Canada Discovery Grant (J.E.H.). P.B. is supported by a Devolved Scholarship from the University of Saskatchewan. The authors would like to thank Champika Fernando for technical support.

---

**Supplemental Figure 1:**
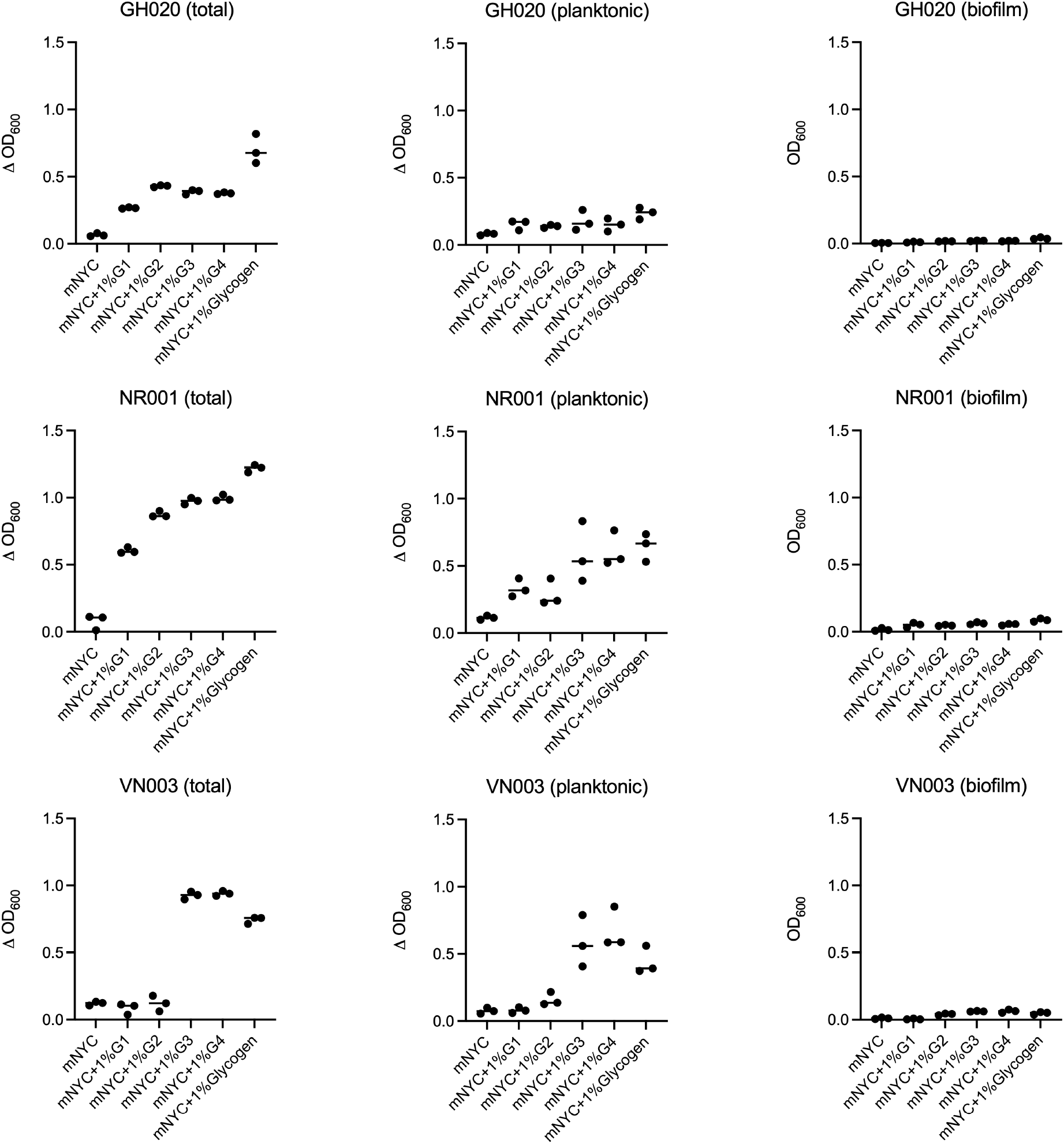
Measurement of total (left), planktonic (middle) and biofilm (right) growth for selected isolates of *G. piotii* (GH020), *G. vaginalis* (NR001) and *G. leopoldii* (VN003). Data from three technical replicates are shown; horizontal line indicates the median. Total and planktonic growth are shown as the increase in OD600 from 0-48h. Biofilm growth was measured using a crystal violet stain as described in the text.

